# Evolutionary Divergence of Structure-Function Coupling between Human and Macaque: Spatial Patterns and Transcriptomic Basis

**DOI:** 10.64898/2026.03.02.708906

**Authors:** Jianing Ma, Wei Li, Yingzi Ma, Jin Chen, Jing Su, Yashi Wu, Chengsi Luo, Wen Li, Jiaojian Wang

**Affiliations:** State Key Laboratory of Primate Biomedical Research, Institute of Primate Translational Medicine, Kunming University of Science and Technology, Kunming 650500, China; Yunnan Key Laboratory of Primate Biomedical Research, Kunming 650500, China; Brainnetome Center, Institute of Automation, Chinese Academy of Sciences, Beijing 100190, China

**Keywords:** structural-functional coupling, cross-species comparison, evolution, molecular mechanisms

## Abstract

Structural-functional coupling (SFC) provides critical insights into how the brain’s structural architecture constrains functional organization for supporting higher-order cognition. Investigating evolutionary differences in SFC between humans and macaques may provide new insights into the neural basis of unique human cognitive abilities. In this study, we analyzed multimodal magnetic resonance imaging data from anesthetized and awake adult rhesus macaques and adult humans to examine cross-species divergences in SFC. Moreover, we integrated transcriptomic data to elucidate the molecular mechanisms underlying evolutionary differences in SFC patterns. We find that humans and macaques exhibit distinct SFC patterns: the human brain shows high SFC in the lateral and medial prefrontal cortex, whereas macaques show high SFC in the sensorimotor cortex. Notably, language-related regions in the human lateral temporal cortex exhibit relatively low SFC. Furthermore, the human whole-brain SFC pattern and the evolutionary differences in SFC between humans and macaques are negatively correlated with cortical evolutionary expansion. By integrating human and macaque transcriptomes, we reveal that the macaque SFC specifically associated genes are primarily involved in basic physiological functions, whereas the human SFC specifically associated genes exhibit evolutionary adaptations in synaptic function, neurotransmitter secretion, and other molecular processes. Moreover, the human-specific genes showing significant overlap with Human Accelerated Regions genes were mainly enriched in cell types of astrocyte and oligodendrocyte and in diseases of schizophrenia and Alzheimer’s diseases. Overall, these findings advance our understanding of the intricate relationships of SFC in human and macaque brains and provide novel insights in understanding evolutionary conservation and species specificity in cognitive function and gene regulation.

## Introduction

Brain structure provides the scaffolding for intrinsic function, with functional regions constrained by the underlying white matter network (1). The brain structural-functional coupling (SFC) describes the complex interplay between anatomical architecture and functional dynamics, revealing hierarchical patterns that link functional brain activity to structural networks (2–4). At the macroscale, SFC varies significantly across different cortical regions and is closely linked to individual cognitive abilities and genetic factors (2, 5, 6). At the microscale, the heterogeneous development of SFC is closely linked to the expression profiles of genes involved in oligodendrocyte- and astrocyte-related pathways (7). This association offers new insights into the fundamental processes underlying brain development, cognitive function maturation, and functional deficits in diseases.

SFC is a particularly useful approach to investigate brain evolution since it reflects both the functional complexity and the cortical refinement during evolutionary expansion (5, 8). SFC provides an alternative way to investigate the origins of higher-order cognitive functions, such as language, decision making, social cognition, and their underlying neural adaptation mechanisms in humans (9, 10). Non-human primates, especially the macaque, which show high similarity in genetics, physiology, and brain structure with humans, are one of the most suitable proxies to explore the neurobiological mechanisms underlying normal cognitive functions and brain disorders (11–13). Although structural and functional differences between humans and macaques have been reported (14–16), it remains unclear how the structural connectome shapes functional connectivity in humans and macaques to promote the brain specializations unique to humans.

Cortical SFC exhibits strong heritability (17) and aligns closely with genetically determined connectivity profiles (18), indicating that its formation is under genetic regulation. Brain development and evolution are widely modulated by spatiotemporal patterns of gene expression (19, 20). This genetic influence implies that mapping SFC onto spatial gene-expression profiles could uncover molecular pathways that shape brain structure-function relationships. The Allen Human Brain Atlas (AHBA) (21) serves as a comprehensive human transcriptomic reference, facilitating the identification of genes that covary with neuroimaging phenotypes and enabling downstream analyses such as pathway and cell-type enrichment (22–24). Recently, a complementary macaque whole-brain transcriptomic atlas is available (25), permitting to explore the shared and evolution-specific molecular basis underlying cross-species same neuroimaging phenotypes. In addition, SFC is also shaped by specific cell types, particularly the glia cells through regulating its key underpinnings of intracortical myelination and excitation and inhibition (E/I) for efficient information processing and higher-order cognition (7, 26–28). Thus, integrating the human and macaque whole brain transcriptome to reveal the molecular basis underlying SFC evolution may shed new light on the origin of human higher-order cognitive functions.

In this study, we aimed to investigate the evolutionary differences in SFC between humans and macaques and to explore their underlying molecular basis by integrating multimodal magnetic resonance imaging (MRI) data from 57 adult rhesus macaques (ages 6-10 years) and 62 adult humans (ages 19-28 years) acquired using the same scanner in a single site and transcriptomic datasets. First, we mapped structural and functional connectomes and constructed an individual SFC matrix for each subject in both species. Second, we compared the macroscopic spatial distributions of SFC across species and identified regions exhibiting significant evolutionary expansion. In addition, we validated the macaque SFC patterns derived from anesthetized data using an independent awake-macaque dataset. Third, we performed spatial association analyses linking SFC patterns to transcriptomic data to identify shared and species-specific genes associated with SFC patterns, and Gene Ontology (GO) enrichment, Kyoto Encyclopedia of Genes and Genomes (KEGG) pathway analyses, and protein-protein interaction (PPI) network analyses were employed to elucidate their molecular functions. We also linked the human-specific genes to cell types and diseases to reveal which cell types regulating human SFC and to identify the evolutionary associations with brain disorders. Finally, we identified genes associated with Human Accelerated Regions (HARs) that are significantly correlated with SFC, and characterized their principal functional roles. A detailed schematic of the study workflow is provided in Figure 1.

**Figure 1.**
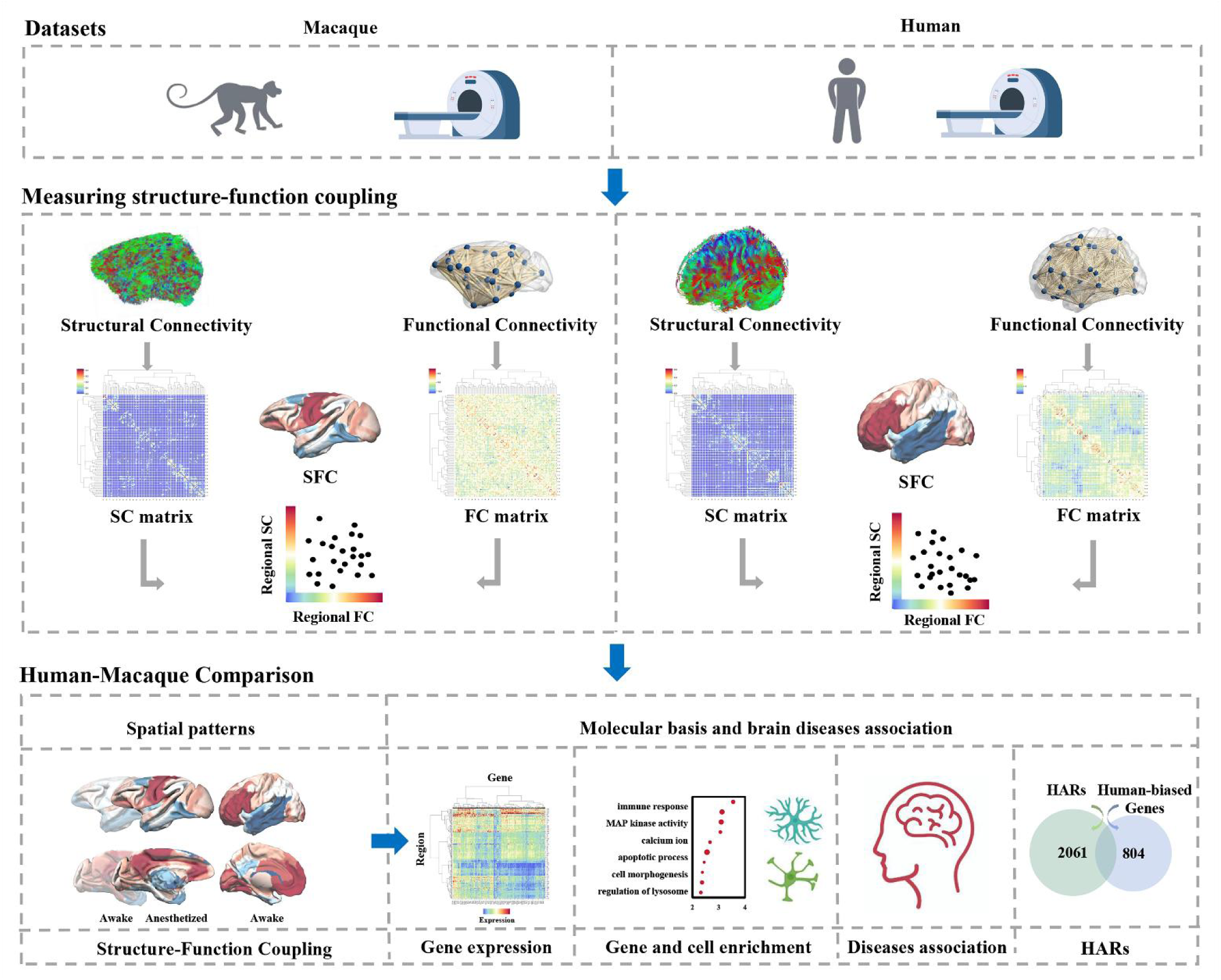
Overview of the methodology pipeline. First, individual structure-function coupling (SFC) matrices were constructed by calculating Spearman’s rank correlations between the non-zero elements of regional structural and functional connectivity profiles. Cross-species comparisons were then performed to identify regions exhibiting significant differences in SFC. Spatial association analyses were conducted to link cortical SFC organization with transcriptomic architecture, followed by GO, KEGG, PPI, cell types, and brain diseases enrichment analyses to elucidate the molecular mechanisms underlying SFC organization and its evolutionary association with brain disorders. In addition, HAR-associated genes that showed significant correlations with SFC were identified.

## Materials and Methods

### Macaques and MRI data acquisition

A total of 54 rhesus macaques with 57 scans (38 females, mean age = 8.11 ± 1.44 years, age range: 6-10 years) from the State Key Laboratory of Primate Biomedical Research, Institute of Primate Translational Medicine, Kunming University of Science and Technology were used in this study (Table 1). All experimental procedures were conducted in strict compliance with national guidelines and regulations on animal protection and use, and the protocols were approved by the National Bureau of Animal Research of China and the Animal Care Committee (permit number: KUST202301018). MRI data were acquired using a Siemens 3T Prisma. Before scanning, the animals were intramuscularly pre-anesthetized with ketamine hydrochloride (15 mg/kg; Gutian, China) and xylazine (1.5 mg/kg; Huamu, China). The anesthesia was stable for at least 1 hour before the MRI acquisition. The scanning parameters are as follows: Diffusion tensor imaging: 30 directions with b = 1000 s/mm², 1 non-diffusion-weighted image with b = 0 s/mm², repetition time (TR)/echo time (TE) = 8500/77 ms, flip angle = 90°, field of view (FOV) = 128 × 128 mm², matrix size = 128 × 128, voxel size = 1 × 1 × 1 mm³, 30 axial slices. Resting-state fMRI: TR/TE = 2000/22 ms, flip angle = 90°, 30 slices, slice thickness = 1.5 mm, voxel size = 1.5 × 1.5 × 1.5 mm³, matrix size = 64 × 64, FOV = 96 × 96 mm², 250 time points.

**Table 1:** Demographics and scanning information of human participants and rhesus macaques.

| Species | Site | Scanning state | Number (M/F) | Mean Age ± SD (years) | Age range (years) |
| --- | --- | --- | --- | --- | --- |
| Humans | KUST | Awake | 24/38 | $22.06 \pm 2.15$ | 19-28 |
| Rhesus macaques | KUST | Anesthetized | 19/38 | $8.11 \pm 1.44$ | 6-10 |
| Rhesus macaques | Newcastle University | Awake | 7/0 | $9.27 \pm 2.09$ | 6-13 |

KUST: Kunming University of Science and Technology; Newcastle University: University of Newcastle upon Tyne.

Given the potential influence of anesthesia on functional connectivity analysis (29), we conducted validation analyses using an independent, publicly available awake macaque resting-state fMRI dataset (30). The awake macaque dataset (Newcastle dataset) comprised 14 rhesus macaques (*Macaca mulatta*) scanned without contrast agents. After strict evaluation of data integrity and motion-related quality control, 7 male animals (mean age = 9.27 ± 2.09 years, age range: 6-13 years) were retained for analysis. All the experimental protocols were approved by the UK Home Office and conducted in accordance with the Animals (Scientific Procedures) Act of 1986 and the European Directive on the Protection of Animals Used for Scientific Purposes (2010/63/EU) (see (31) for detailed procedures on awake imaging preparation). Imaging was scanned on a Vertical Bruker 4.7T primate-dedicated scanner in awake, head-fixed animals across two sessions, with acquisition parameters of TR = 2600 ms, TE = 17 ms, and a voxel size of 1.22 × 1.22 × 1.24 mm³. Data were downloaded from the NHP data-sharing consortium PRIME-DE (http://fcon_1000.projects.nitrc.org/indi/indiPRIME.html) (32).

### Human subjects and MRI data acquisition

The human MRI data used in this study were obtained from the National Key Laboratory of Primate Biomedical Research, Institute of Primate Translational Medicine, Kunming University of Science and Technology. All participants provided written informed consent prior to their inclusion in the study. After rigorous assessment of data completeness and quality control, a total of 62 healthy participants (38 females; mean age = 22.05 ± 2.15 years; age range: 19-28 years) were included in the final analysis (permit number: KMUST-MEC-207) (Table 1). All participants were scanned using a Siemens 3T 64-channel MRI scanner. Before scanning, participants were asked to relax and remain still with their eyes opened, avoiding falling asleep. The scanning parameters for data acquisition are as follows: Diffusion tensor imaging: 64 directions with b = 1000 s/mm², 1 non-diffusion-weighted image with b = 0 s/mm², TR/TE = 3900/72 ms, 70 axial slices, FOV = 256 × 256 mm², matrix size = 128 × 128, voxel size = 2 × 2 × 2 mm³. Resting-state fMRI: TR/TE = 1500/30 ms, flip angle = 70°, 40 slices, slice thickness = 3 mm, voxel size = 3.0 × 3.0 × 3.0 mm³, matrix size = 64 × 64, FOV = 192 × 192 mm², 250 time points.

### Human and macaque diffusion MRI data preprocessing and anatomical network mapping

FSL software (http://www.fmrib.ox.ac.uk/fsl) was used for diffusion MRI data processing (33, 34). First, head motion and eddy current correction were performed using the eddy-correct tool in FSL to mitigate scan-related artifacts. Diffusion gradient vectors were also rotated to adjust for the subject’s estimated motion (35). We applied the ball-and-sticks diffusion model to each subject’s DTI data to estimate fiber orientation using the FSL bedpost function, which employs Markov chain Monte Carlo sampling to estimate principal fiber orientation and diffusion characteristics per voxel (36, 37). Subsequently, probabilistic tractography was conducted through the FSL probtrackx function, which iteratively samples voxel-level fiber orientation distributions to map the spatial pathways and connectivity strength between predefined seed and target regions (37–40). The connectivity probability was calculated as the total of streamlines in target brain region divided by the total sampling streamlines of the seed region (5000 × total number of voxels). Given the fiber tracking results affected by the seeding locations, the connectivity probability from seed to target regions is usually not equivalent to the connectivity probability from target to seed regions. Thus, the connectivity probability between the seed area and the target area is defined by averaging the two connectivity values. The same methods and procedures were used for diffusion MRI data preprocessing and anatomical connectivity mapping for both human and macaque subjects. Using the cross-species Regional Map (RM) atlas, the brains of humans and macaques were parcellated into 92 subregions, and 92 × 92 anatomical connectivity matrices were subsequently constructed for each subject (41).

### Human and macaque resting-state fMRI data preprocessing and functional network mapping

The human resting-state fMRI data were preprocessed as follows: first, the initial 10 time points were discarded and realigned to correct head motion. Next, the data were normalized to the MNI template and resampled to 3 × 3 × 3 mm³. A 6-mm Gaussian kernel was applied to the data for smoothing, and a temporal band-pass filter (0.01-0.1 Hz) was applied. Friston’s 24 motion parameters, white matter, global signal, and cerebrospinal fluid signals were regressed out. Finally, the volumes with frame displacement (FD) > 0.2 mm were scrubbed using cubic spline interpolation to minimize head motion effects. The macaque resting-state fMRI data were preprocessed in the same way as the human data. The first 10 volumes were discarded and realigned to the first volume to correct head motion. Then, all the volumes were spatially normalized to a standard INIA19 template and resampled to 1 × 1 × 1 mm³. A 3-mm Gaussian kernel was applied for smoothing, and a band-pass filter (0.01-0.05 Hz) was used to remove unwanted frequency components. The Friston’s 24 head motion parameters, white matter, cerebrospinal fluid, and mean global signals were regressed out. To further exclude the head motion effects on functional connectivity analysis, a scrubbing method using cubic spline interpolation was conducted to eliminate frame-wise head motion when the mean frame displacement (FD) > 0.2 mm. After preprocessing, the BOLD time series were averaged within each cortical region defined by the RM atlas (41) which parcellate both human and macaque brains into 92 homologous regions. Pairwise Pearson’s correlations were then computed between all regional time series, and the resulting correlation matrices were transformed using Fisher’s *r*-to-*z* transformation to generate individual functional connectivity (FC) matrices for both species.

### Structure-function coupling analysis

For each participant, the structural and functional connectivity features of each region with all other brain regions were extracted and vectorized. The strength of structure-function coupling (SFC) was assessed by calculating Spearman’s rank correlation coefficient between the structural and functional connectivity features of each region (42, 43). The same procedures were performed for all the brain regions, and a SFC matrix was obtained for each participant.

To characterize the spatial distribution of human-macaque SFC patterns, we employed the Cole-Anticevic Brain-wide Network Partition (CAB-NP) which divides all brain regions into 12 finer-grained networks and includes a specific language network (https://balsa.wustl.edu/study/wZML). The 12 networks include VIS (primary/secondary visual), SMN (somatomotor), DAN (dorsal attention), VMM (ventral multimodal), OAN (orbito-affective), FPN (frontoparietal), DMN (default mode), CON (cingulo-opercular), AUD (auditory), LAN (language), and PMM (posterior multimodal) (44). Using the macaque-to-human spherical deformation field developed by Xu et al. (45), we transformed the human CAB-NP 12 network labels into the macaque FSL 32k surface, thereby generating a 12 network partition for the macaque brain. We averaged the regional SFC values within each of the 12 CAB-NP networks to obtain network-level SFC distribution patterns for both species. In addition, to verify whether the average SFC patterns derived from the CAB-NP parcellation were consistent with those found based on the traditional Yeo’s 7 functional networks, we further parcellated the brain into 7 canonical functional networks, including the default mode network (DMN), frontoparietal network (FPN), dorsal attention network (DOR), ventral attention network (VEN), somatosensory network (SOM), limbic network (LIM), and visual network (VIS) (46, 47). Each brain region was assigned to one of these networks, and the mean SFC within each network was computed separately for humans and macaques.

### Validation analysis using deterministic tractography

To validate the analytical results, we also constructed structural brain network mapping using deterministic tractography (48). Deterministic tractography using the fiber assignment by continuous tracking (FACT) method was conducted with DiffusionKit software (https://diffusionkit.readthedocs.io/en/latest/) (49). The tractography terminated when the curvature angle threshold exceeded 55 degrees or the fractional anisotropy (FA) dropped below 0.1. The step size of streamlines was set at 0.5 mm. The structural connectivity strength was quantified by counting the number of fiber bundles connecting each pair of regions, and the mean FA, mean diffusivity (MD) and number of streamlines (NUM) values were calculated to define the weighted anatomical connectivity network. After obtaining the anatomical connectivity matrix for each subject, the SFC matrix was also calculated using Spearman’s rank correlation. The similarity between SFC patterns derived from the probabilistic and deterministic tracking methods were quantified using Pearson’s correlation coefficient.

### Validation analysis using awake macaque’s fMRI data

Although a previous study indicated that the spatial distribution of functional flexibility in the anesthetized macaque brain is significantly correlated with that in the awake human brain (50), we still further analyzed publicly available resting-state fMRI data collected under awake conditions to evaluate the potential influence of anesthesia on our findings. We applied identical preprocessing procedures to the resting-state fMRI data from both anesthetized and awake macaques. A Pearson correlation was performed between the whole-brain mean FC in all macaques under awake and anesthetized conditions to evaluate the global similarity. Next, we evaluated the SFC of macaques using three complementary approaches: (1) Regional group-level SFC based on the correlation between group-averaged SC and group-level average awake FC matrices; (2) Regional group-level SFC based on the correlation between group-averaged SC and group-level average anesthetized FC matrices; (3) Regional individual-level SFC calculated by correlating SC and FC matrices for each subject (both under anesthesia) and averaging the resulting SFC maps across subjects. Finally, pairwise Pearson correlations were performed among the three resulting measures to assess the similarity of SFC patterns.

### Evolutionary areal expansion and structure-function coupling

Evolutionary cortical surface area expansion between humans and macaques was estimated by quantifying the degree of surface deformation required to spatially align human cortical regions with their macaque homologues. These expansion estimates were derived from a publicly available atlas (45). Higher expansion values indicate regions that have undergone greater evolutionary enlargement. To investigate whether regions that underwent different degrees of expansion during evolution also exhibit distinct SFC pattern, we assessed the spatial correspondence between the evolutionary expansion map and the mean human cortical SFC distribution. The statistical significance of this spatial correlation was evaluated using a conservative spatial permutation test, which generates a null distribution of randomly rotated brain maps while preserving the intrinsic spatial autocorrelation structure of the cortical surface (51). Specifically, the mean SFC map was projected onto an fsLR-32k spherical surface and randomly rotated 10,000 times to produce null maps that preserve spatial neighborhood relationships. For each rotation, regional values were extracted as the mode of vertices within each cortical parcel, and Pearson’s correlation coefficients between the rotated SFC values and the corresponding evolutionary expansion values were computed to generate a null distribution. The permutation-based *p*-value was defined as the proportion of null correlations exceeding the observed empirical correlation, allowing us to determine whether the spatial association between cortical expansion and SFC deviated significantly from chance. To further investigate the evolutionary relevance of SFC, we examined whether regional differences in SFC between humans and macaques were spatially aligned with interspecies cortical expansion. Regional SFC values were compared between species using two-sample t-tests with Bonferroni correction for multiple comparisons (*p* < 0.05). The top four regions with the highest and lowest t-values were displayed for visualization. We then computed the Pearson correlation between the whole-brain human-macaque SFC difference map and the evolutionary areal expansion map reported by Xu et al. Statistical significance was again assessed using the same spatial permutation (“spin test”) approach to ensure that the observed correlation was not confounded by spatial autocorrelation across the cortical surface.

### Functional decoding of whole-brain SFC pattern

To reveal cognitive functions primarily associated with the whole brain average SFC across all subjects, we performed functional decoding using the NeuroSynth meta-analytic framework. Functional associations linked to SFC were examined following the 24 topic terms identified by Margulies et al. (52). Twenty binary masks were created by segmenting the whole brain average SFC map values into 20 bins with 5% increments. Terms were ranked by the weighted average of *z*-statistics for visualization purposes, and relevant cognitive functions were identified through FDR correction (*p* < 0.05).

### Transcription-neuroimaging association analysis

To explore the molecular basis underlying evolutionary differences in SFC patterns, spatial correlation between transcriptomic expression profiles and SFC pattern was assessed. Human gene expression data were accessed from the Allen Human Brain Atlas (AHBA) with samples from six post-mortem donors (5 males, 1 female), aged 24-57 years (mean age 42.5 ± 13.4) (https://human.brain-map.org/). The AHBA microarray expression data were preprocessed and integrated into a matrix of 92 regions of interest (ROIs) × 15,637 genes (21, 53, 54). We only used the data from the left hemisphere to reduce variability across regions and hemispheres due to sample availability differences (53). Macaque gene expression data from the brain-wide and cell-specific transcriptomic atlas was created by integrating bulk RNA sequencing datasets and single nucleus RNA sequencing datasets from 819 samples from 110 brain regions (8 males and 1 female; mean age = 13.6 ± 7.8 years) (25). To ensure cross-species comparability, we selected only left-hemisphere regions, yielding 46 homologous areas shared by humans and macaques. We ultimately selected 11,033 genes with matching names between humans and macaques to construct a 46 × 11,033 gene expression matrix (46 left-hemisphere homologous regions, 11,033 genes) for subsequent analyses. Finally, the spatial correlations between the genes’ expression profiles and the SFC patterns were calculated to identify the related genes.

Partial least squares (PLS) regression was used to investigate the molecular basis of SFC. For humans and macaques, SFC patterns in the left cortical regions served as the response variable, while the corresponding regional gene expression profiles were used as predictor variables. The significance of PLS components was tested using a permutation approach, where the response variable was randomly permuted 1,000 times to calculate a *p*-value for each component. PLS gene weights were normalized to *z*-scores by dividing each weight by the standard deviation derived from bootstrapping, enabling the ranking of genes. Significantly associated genes with SFC were identified from the PLS results using an FDR-corrected threshold of *p* < 0.05. The top five positively and negatively ranked genes based on their *z*-scores were selected separately for humans and macaques to visualize gene expression in the left hemisphere. The genes that were significantly associated only in humans were labeled as human-biased, those only in macaques as macaque-biased, and those significant in both species as shared genes.

### Transcriptomic characterization and functional enrichment Analysis

To characterize the spatial organization and functional relevance of species-biased genes, we first mapped the average expression patterns of the human-biased and macaque-biased gene sets and quantified their distributions across the 12 CAB-NP functional networks in left-hemisphere. In addition, functional decoding was also performed for the left-hemispheric gene expression map significantly associated with human SFC as described above to reveal the cognitive domains linked to the spatial expression pattern of human-specific genes associated with SFC providing a complementary insight into how transcriptomic variations contribute to higher-order cognitive functions. For cross-species comparability, Gene Ontology (GO) and Kyoto Encyclopedia of Genes and Genomes (KEGG) enrichment analyses were performed using the R package clusterProfiler to identify enriched biological processes, molecular functions, cellular components, and signaling pathways. Significance was assessed using the FDR correction with a threshold of *p* < 0.05. Enrichment analyses for human-biased genes were performed using the org.Hs.eg.db library from the OrgDb database, and enrichment for macaque-biased genes was conducted using the org.Macaca (AH114564) package.

### Cell-type specificity and disease enrichment analyses

To assess whether the genes specifically associated with human SFC show cell type-specific expression, we first assigned these genes to 58 cell-type labels derived from five single-cell studies (55–59). These labels were then grouped into eight canonical classes (excitatory neurons, inhibitory neurons, astrocytes, microglia, endothelial cells, oligodendrocytes, oligodendrocyte precursor cells, and synapse-related genes) following the annotation framework of Seidlitz et al (60). Overlap between the target gene set and each cell-type list was evaluated via permutation testing with 5,000 random resampling to generate null distributions, and resulting *p* values were adjusted for multiple comparisons using FDR correction (*p* < 0.05) to identify significant enrichments. In addition, we conducted an exploratory analysis to examine whether the human-biased genes associated with human neuropsychiatric disorders to explore the evolutionary processes on human brain disorders. We performed disease enrichment analysis on the identified human-specific genes using the ToppGene Suite (https://toppgene.cchmc.org/). Briefly, the human-specific gene list was uploaded to ToppFun, where enrichment significance was evaluated using the hypergeometric test (equivalent to a one-sided Fisher’s exact test (61)). The resulting *p*-values were adjusted for multiple comparisons using the FDR correction. Only disease terms with FDR < 0.05 were considered significantly enriched.

### PPI network and functional enrichment of overlapping genes

We additionally evaluated conserved molecular interactions among the overlapping genes, we performed protein-protein interaction (PPI) analysis using STRING v11.0 (https://cn.string-db.org/), with a maximum confidence interaction score of 0.7, and the genes with the top 10% degree values in the network were identified as hub genes. The whole-brain expression patterns of the top three hub genes, ranked by node degree, were visualized. Finally, GO and KEGG enrichment analyses were conducted on the shared gene set to highlight conserved biological pathways.

### HAR-associated genes

To determine whether the human-biased genes associated with human SFC were localized with the human accelerated regions (HAR) genes, the overlap genes were identified. The HAR genes were accessed from a previous study (62). The 2,162 HAR genes were compared with the 905 human-specific genes significantly associated with SFC, yielding 101 overlapping genes. To further characterize these overlapping genes, we first mapped the spatial expression patterns of the 101 genes across cortical regions to visualize their distribution in the left hemisphere. Next, we quantified the mean expression of these genes within the 12 canonical functional networks defined by the CAB-NP to assess network-level distribution. Gene set enrichment analyses were then performed using the GO and KEGG databases to identify biological functions and signaling pathways related to these 101 genes. Finally, we conducted functional decoding of the cortical expression map using the NeuroSynth meta-analytic framework to link spatial expression patterns with 24 cognitive domains.

## Results

### Evolutionary differences in spatial patterns of cortical SFC between human and macaque

Using spatial Spearman’s rank correlation, the spatial distribution patterns of SFC in humans and macaques and significant regional differences were identified. In macaques, high SFC was observed in the somatosensory cortex and visual cortex while relatively low SFC was observed in the dorsolateral prefrontal cortex (dlPFC) (Fig. 2A). In humans, high SFC was observed in the prefrontal cortex (PFC) and primary visual cortex, while low levels of SFC were detected in the lateral temporal regions (Fig. 2B). To characterize whether the spatial distribution of SFC is different in language network between human and macaque, we applied the CAB-NP to partition all brain regions into 12 functional networks (Fig. S1A). In macaques, the sensorimotor (SMN) and primary visual (VIS1) demonstrated the highest SFC while the default mode (DMN) and ventral multimodal (VMM) exhibited the lowest SFC (Fig. 2C). In humans, the orbito-affective (OAN) and VIS1 exhibited the highest SFC, whereas the language (LAN) and VMM showed the lowest SFC (Fig. 2D). In addition, we also partitioned the brain into 7 functional networks and found similar functional distribution (Fig. S1B). The macaque showed high SFC in SOM and ventral attention network (VEN) while human exhibited high SFC in FPN and VIS (Fig. S1C, Fig. S1D). We further examined the spatial relationship of human SFC patterns with cortical expansion and found a significant negative correlation between the whole-brain SFC distribution and the evolutionary areal expansion map, indicating that regions undergoing greater evolutionary expansion tend to exhibit weaker SFC in humans (Fig. 2E). To further assess the evolutionary relevance of SFC, we observed a significant negative spatial correlation between the interspecies SFC difference map (human-macaque t-values) and cortical areal expansion (Fig. 2F). The interspecies SFC difference map showed the most pronounced difference in the medial prefrontal cortex, whereas the smallest difference in the superior temporal cortex (Fig. S2A, Fig.S2B). Our analysis further identified the behavioral and cognitive functions associated with human SFC pattern, providing a heatmap that displays the weighted mean z-statistics of cognitive components, sorted by rows. Brain regions with lower SFC values in humans were primarily associated with emotion and declarative memory, whereas regions with higher human SFC values were linked to motor and visuospatial (Fig. S3).

**Figure 2.**
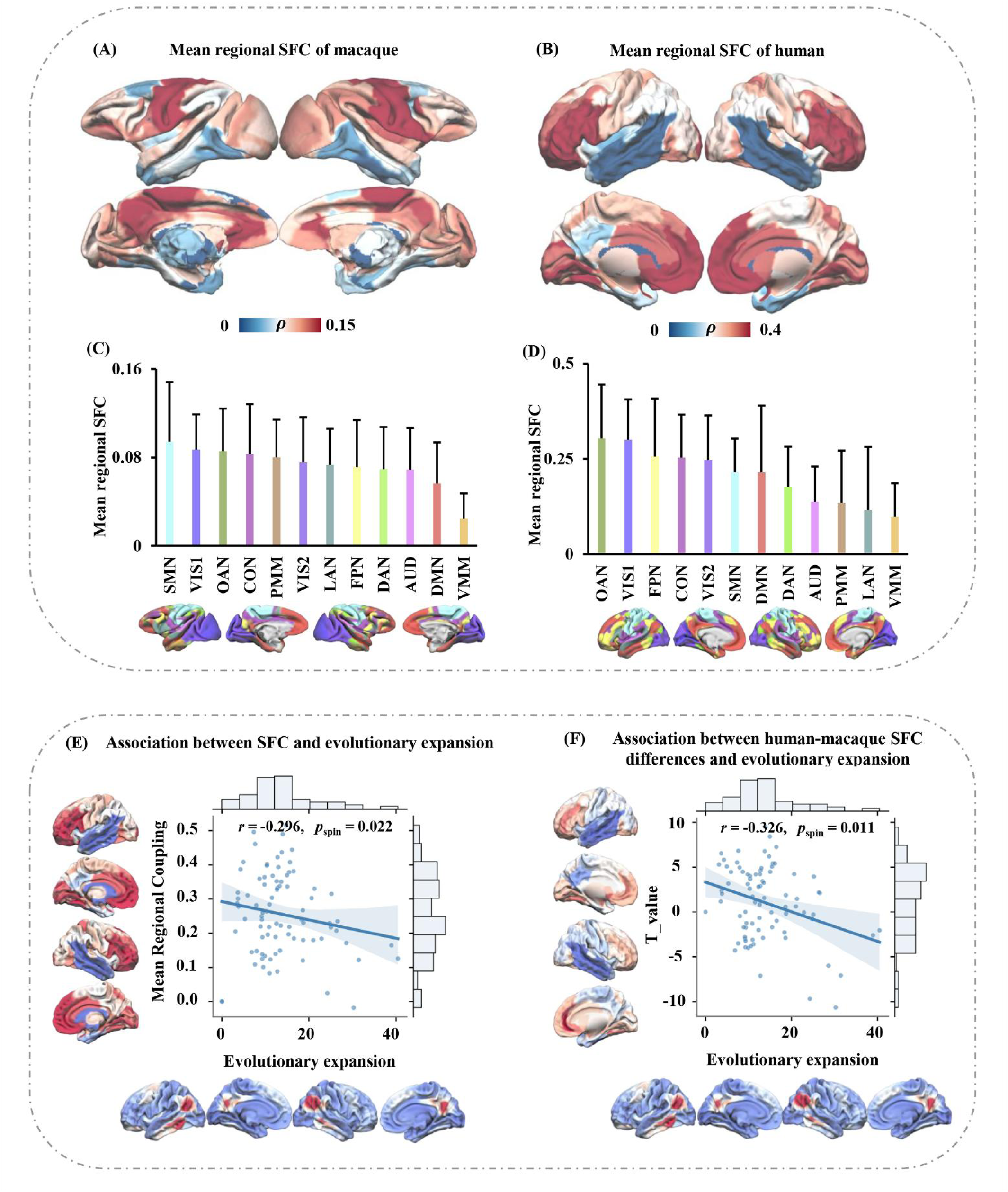
Macroscale distribution of SFC and association with evolutionary expansion. Spatial distribution patterns of SFC in macaque (A) and human brains (B). SFC distributions in humans (C) and macaques (D) were examined across the 12 functional networks defined by the CAB-NP. (E) Spatial permutation ("spin") tests revealed a significant negative correlation between human cortical SFC and evolutionary areal expansion. (F) A similar negative association was observed between interspecies SFC differences (human-macaque t-values) and cortical expansion. Note: VIS, primary/secondary visual; SMN, somatomotor; DAN, dorsal attention; VMM, ventral multimodal; OAN, orbito-affective; FPN, frontoparietal; DMN, default mode; CON, cingulo-opercular; AUD, auditory; LAN, language; PMM, posterior multimodal. *p_spin_*: spin test.

To validate the robustness of our findings, we further used a deterministic fiber tracking method to investigate the SFC patterns in both humans and macaques. Spatial correlation analyses indicated that the SFC patterns obtained using the deterministic tracking approach are similar to those obtained using probabilistic tracking in both macaques (probabilistic & number weighted: *r* = 0.75, *p* < 0.001; probabilistic & MD weighted: *r* = 0.46, *p* < 0.001; probabilistic & FA weighted: *r* = 0.23, *p* < 0.005) and humans (probabilistic & number weighted: *r* = 0.65, *p* < 0.001; probabilistic & MD weighted: *r* = 0.56, *p* < 0.001; probabilistic & FA weighted: *r* = 0.41, *p* < 0.001) (Fig. S4, Fig. S5).

To further evaluate the reliability of SFC pattern of anesthetized macaque, we examined the spatial similarity of whole brain SFC patterns derived from anesthetized and awake macaques. A significant positive Pearson correlation was observed between awake and anesthetized whole-brain FC (*r* = 0.66, *p* < 0.001), indicating that the overall FC topology is largely preserved in anesthetized (Fig. 3A). We next assessed the SFC similarities calculated using different approaches. Specifically, the regional group-level SFC defined using group-averaged SC and group-level average awake FC matrices was significantly correlated with the regional group-level SFC derived from group-averaged SC and group-level average anesthetized FC (*r* = 0.59, *p* < 0.001) and with the regional individual-level SFC derived from subject-wise SFC correlations under anesthesia (*r* = 0.45, *p* < 0.001). Moreover, the regional individual-level SFC showed a strong correlation with the regional group-level SFC defined using group-averaged SC and group-level average anesthetized FC (*r* = 0.81, *p* < 0.001) (Fig. 3B). Together, these results demonstrate that both FC and SFC exhibit high similarity in spatial patterns between awake and anesthetized states, supporting the validity of the SFC findings using anesthetized macaque data.

**Figure 3.**
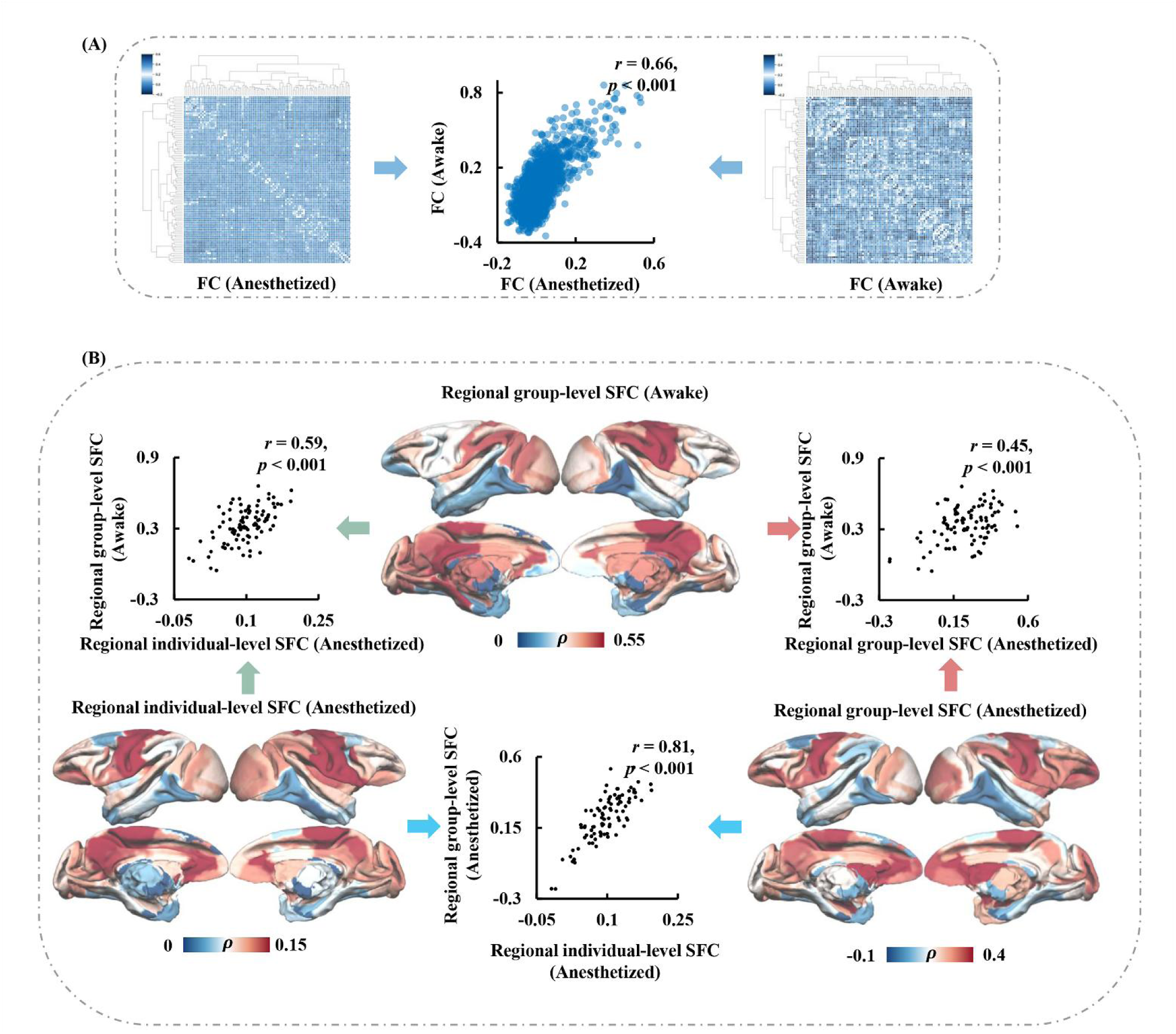
Similarity analyses of functional connectivity (FC) and SFC between awake and anesthetized macaques. (A) Group-averaged whole-brain FC matrices from awake and anesthetized macaques showed a significantly positive correlation (*r* = 0.66, *p* < 0.001), indicating broadly similar large-scale functional organization across states. (B) Three complementary SFC measures were evaluated: (Top) Regional group-level SFC (Awake), computed by Spearman rank correlation between the group-averaged SC and the group-averaged awake FC; (Right) Regional group-level SFC (Anesthetized), computed by Spearman rank correlation between the group-averaged SC and the group-averaged anesthetized FC; and (Left) Regional individual-level SFC (Anesthetized), derived by first computing SFC maps for each anesthetized subject using Spearman rank correlation between individual SC and FC matrices, followed by averaging across subjects. These three SFC measures showed moderate to strong pairwise correlations (*r* = 0.45-0.81, all *p* < 0.001), indicating that the SFC patterns observed under anesthesia closely reflect those measured during the awake state.

### Evolutionary difference in molecular basis underlying SFC between human and macaque

We employed PLS analysis to establish the associations between the spatial patterns of SFC and transcriptome profiles in human and macaque. The first PLS component (PLS1) in macaques accounted for the largest contribution in gene expression scores, with a positive correlation observed between PLS1 scores and the spatial pattern of SFC (Pearson’s *r* = 0.55, *p* < 0.001) (Fig. 4A). In humans, PLS1 also captured the largest contribution to gene expression scores (Pearson’s *r* = 0.56, *p* < 0.001) (Fig. 4B). Significant positively and negatively associated genes were identified from PLS1. The top five positively weighted and top five negatively weighted genes (FDR *p* < 0.05) were identified, and their expression patterns in the left hemisphere were visualized for both humans and macaques (Fig. S6). We combined the significant positive and negative genes (*p* < 0.05, FDR corrected) from the PLS analysis to identify human SFC-specific genes and macaque SFC-specific genes. In total, 905 human SFC-specific genes, 4,880 macaque SFC-specific genes, and 818 overlapping genes were identified (Fig. 4C). The whole-brain mean expression patterns of human- and macaque-specific genes revealed that macaque-specific genes were highly expressed in the PMM and AUD networks, whereas human-specific genes showed higher expression in the VMM and OAN networks (Fig. 4D, 4E).

**Figure 4.**
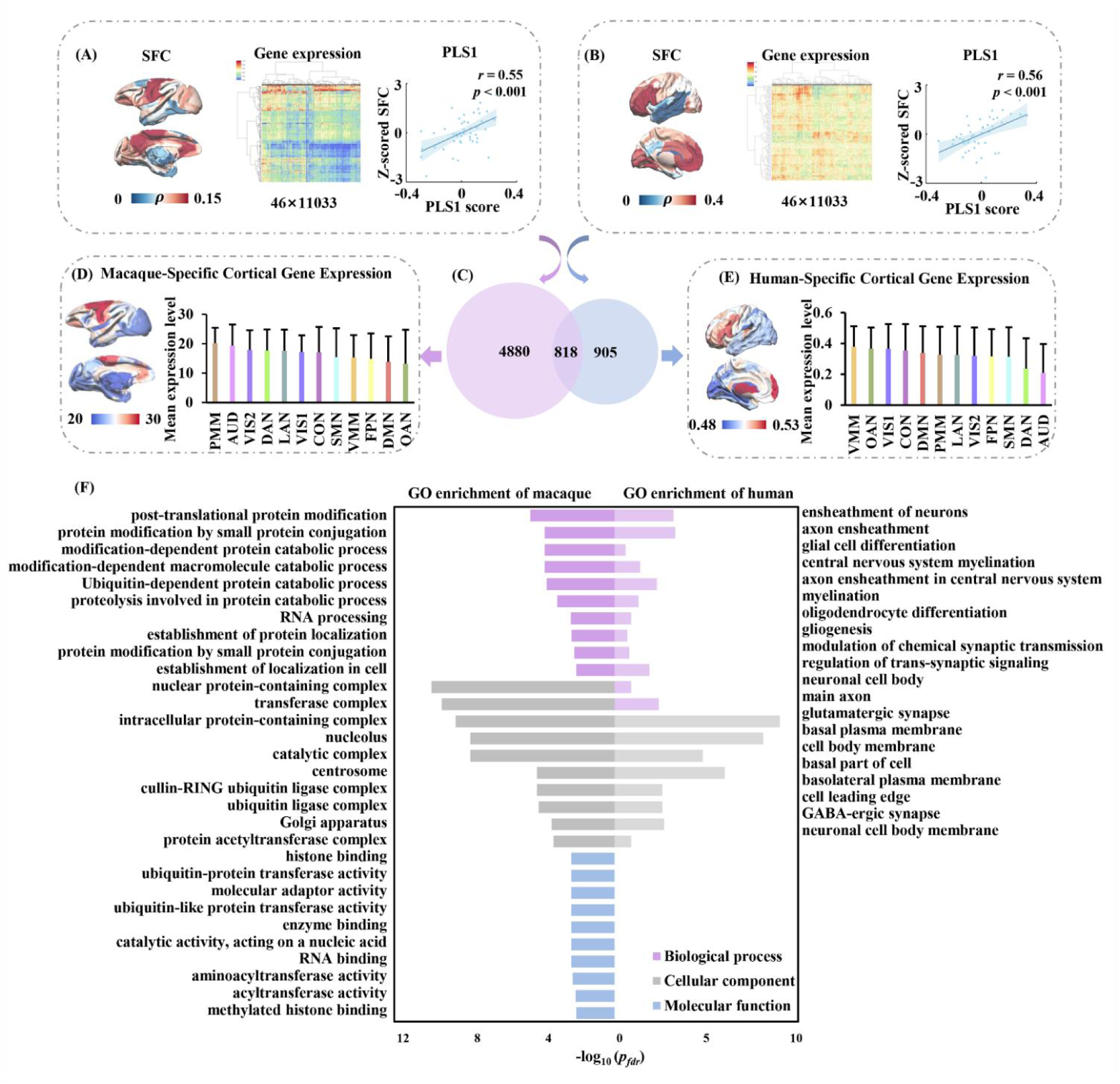
Association between SFC and gene transcription profiles in humans and macaques. (A) Regional SFC values across 46 left-hemisphere homologous areas in macaques (left) and the expression profiles of 11,033 cross-species matched genes derived from the integrated macaque transcriptomic atlas (middle) were shown. The correlation between SFC and the first partial least squares component (PLS1) is presented (right panel). (B) Corresponding analyses in humans, using SFC variations across the same 46 homologous regions (left) and expression profiles of the same 11,033 homologous genes extracted from the Allen Human Brain Atlas (middle). The SFC-PLS1 association is displayed (right panel). (C) Classification of PLS significant genes into macaque-specific (n = 4880), human-specific (n = 905), and shared genes (n = 818). (D, E) Spatial distribution of species-specific gene expression across 12 functional brain networks. (F) GO enrichment analyses of human- and macaque-specific genes, covering Biological Process (BP), Cellular Component (CC), and Molecular Function (MF) categories, highlighting divergent molecular pathways associated with SFC. Note: pfdr: FDR corrected.

Enrichment of macaque-specific genes revealed that these genes are predominantly involved in molecular domains related to proteostasis (post-translational modification and ubiquitin-mediated degradation), intracellular localization and complex assembly (e.g., Golgi apparatus, centrosome), RNA/nucleic-acid processing (e.g., RNA processing, nucleolus) and enzymatic transferase activities (e.g., aminoacyl-/acyltransferases), as well as epigenetic chromatin interactions (e.g., histone binding) (Fig. 4F). Collectively, these pathways underpin core cellular functions including maintaining protein quality, orchestrating gene expression and protein synthesis, organizing subcellular trafficking, and regulating transcriptional programs, which are essential for cellular survival, differentiation and stress adaptation. In contrast, enrichment analysis of human-specific genes revealed a pronounced involvement in neurodevelopmental and cognitive processes. The pathway-level enrichment for these genes further indicated strong involvement in (1) myelination and glial cell differentiation (e.g., axon ensheathment, oligodendrocyte differentiation), supporting efficient signal conduction; (2) synaptic regulation and neuronal projection organization (e.g., modulation of chemical synaptic transmission, regulation of trans-synaptic signaling), reflecting their role in circuit-level information processing; and (3) cellular polarity and membrane compartmentalization (e.g., basal plasma membrane, cell leading edge), which may facilitate axonal guidance and synaptic specificity (Fig. 4F).

To further determine the main cell types regulating human SFC, cell-type enrichment for human-specific genes were conducted and found that these genes were significantly overrepresented in oligodendrocytes, astrocytes, and synapse-related gene categories. This pattern highlights their potential roles in myelination, glial support, and synaptic signaling, respectively (Fig. 5A). In addition, disease enrichment analysis of human-specific genes revealed significant overrepresentation in Schizophrenia, Visual seizure, and Alzheimer’s disease (Fig. 5B). Additionally, we examined the behavioral and cognitive functions related to gene expression of human-specific genes. Functional decoding indicated that the human-specific gene set showed relatively low mean expression in regions associated with visuospatial processing, whereas mean expression was higher in regions linked to reward-based decision making, social cognition, and emotion (Fig. S7).

**Figure 5.**
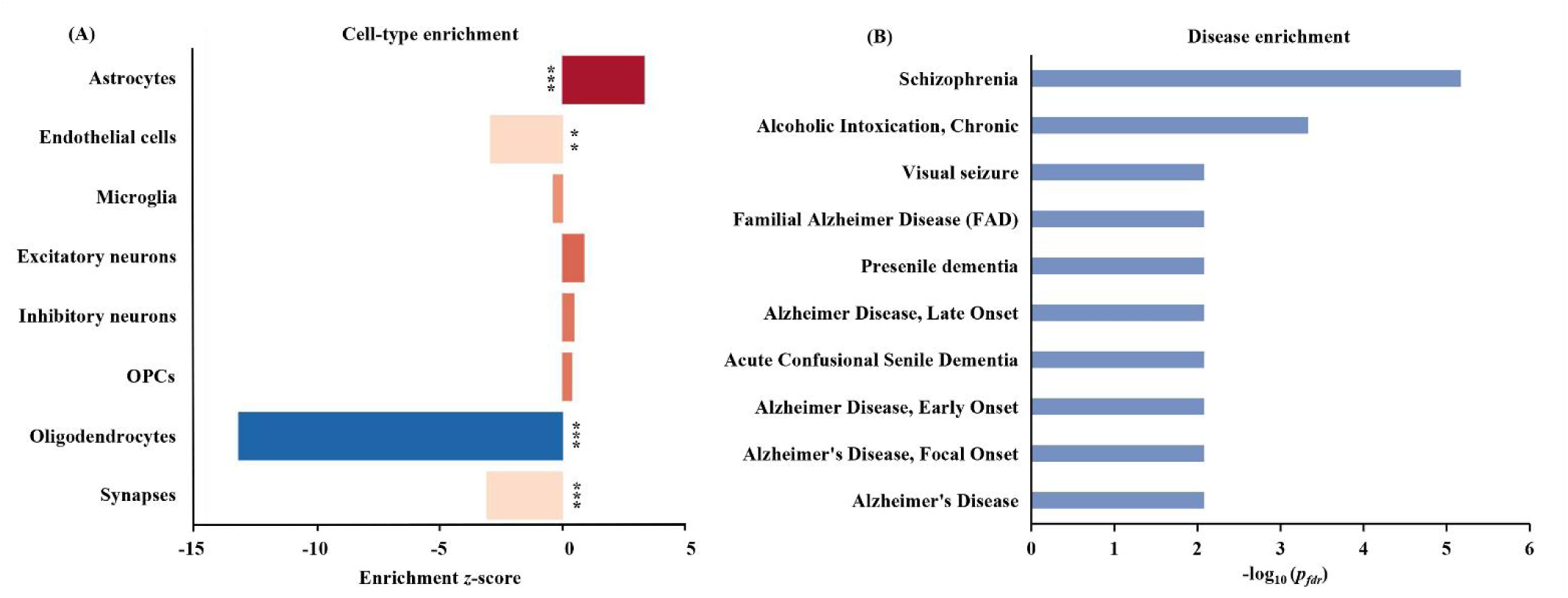
Cell-type and disease enrichment of human-specific genes associated with SFC. (A) Cell-type enrichment analysis showed that human-specific genes were significantly overrepresented in oligodendrocytes, astrocytes, and synapse-related categories, indicating roles in myelination, glial support, and synaptic signaling. (B) Disease enrichment analysis revealed significant associations with Schizophrenia, Visual seizure, and Alzheimer’s disease, based on hypergeometric testing with FDR correction. Note: \*\*\**p* < 0.001; *p_fdr_*: FDR corrected.

Using PPI analysis, we constructed a statistically significant PPI network and identified the top 10 hub genes with the highest degree values within the overlapping gene network (Fig. S8A, Fig. S8B). In addition, we examined the whole-brain expression patterns of the top three hub genes. Notably, the top three hub genes (RPS3A, RPS3, and RACK1) shared by humans and macaques are all housekeeping genes. We also conducted the genes enrichment for the overlapping genes in the human and macaque datasets, respectively. The overlapping genes exhibited broadly similar functional patterns across species, primarily involving fundamental cellular processes required for maintaining cellular homeostasis. Specifically, macaque genes were mainly associated with transcriptional regulation (e.g., RNA polymerase complexes), while human genes were predominantly enriched in protein synthesis and RNA metabolic processes (e.g., translation, ribosome, tRNA processing), as well as cellular structural components (Fig. S8C). Together, these enrichment profiles suggest that, despite small differences, both sets of genes contribute to essential transcriptional and translational functions that underlie basic cellular activity.

### Identifying HAR genes with human-biased genes

To investigate whether the human SFC specifically associated genes were localized in HAR, we compared 2,162 genes located within HARs with 905 human-specific genes significantly associated with SFC. We identified 101 overlapping genes (Fig. 6A). We then examined the spatial expression patterns of these overlapping genes which highly expressed in the prefrontal cortex while lowly expressed in parietal and visual cortices (Fig. 6B). To further characterize their functional roles, we quantified their distribution across 12 functional networks, revealing that these genes highly expressed in networks supporting higher-order cognition, such as VMM and OAN (Fig. 6B). GO analysis of these genes identified significant involvement in biological processes related to synapse organization, axon guidance, and intracellular signal transduction, as well as KEGG pathways associated with neurodevelopment and synaptic plasticity (Fig. 6C). Functional decoding of the cortical expression map derived from the 101 HAR-SFC overlap genes using the NeuroSynth meta-analytic framework revealed systematic associations with distinct cognitive domains (Fig. 6D). The heatmap shows the weighted mean *z*-scores of cognitive terms, ordered by their correspondence with regional gene expression levels. Regions with relatively low expression of the overlap genes were preferentially associated with visuospatial processing and eye-movement functions, whereas regions with relatively high expression were associated with emotion and social cognition, indicating that these HAR-linked, SFC-associated genes are enriched in networks supporting higher-order socio-emotional cognition.

**Figure 6.**
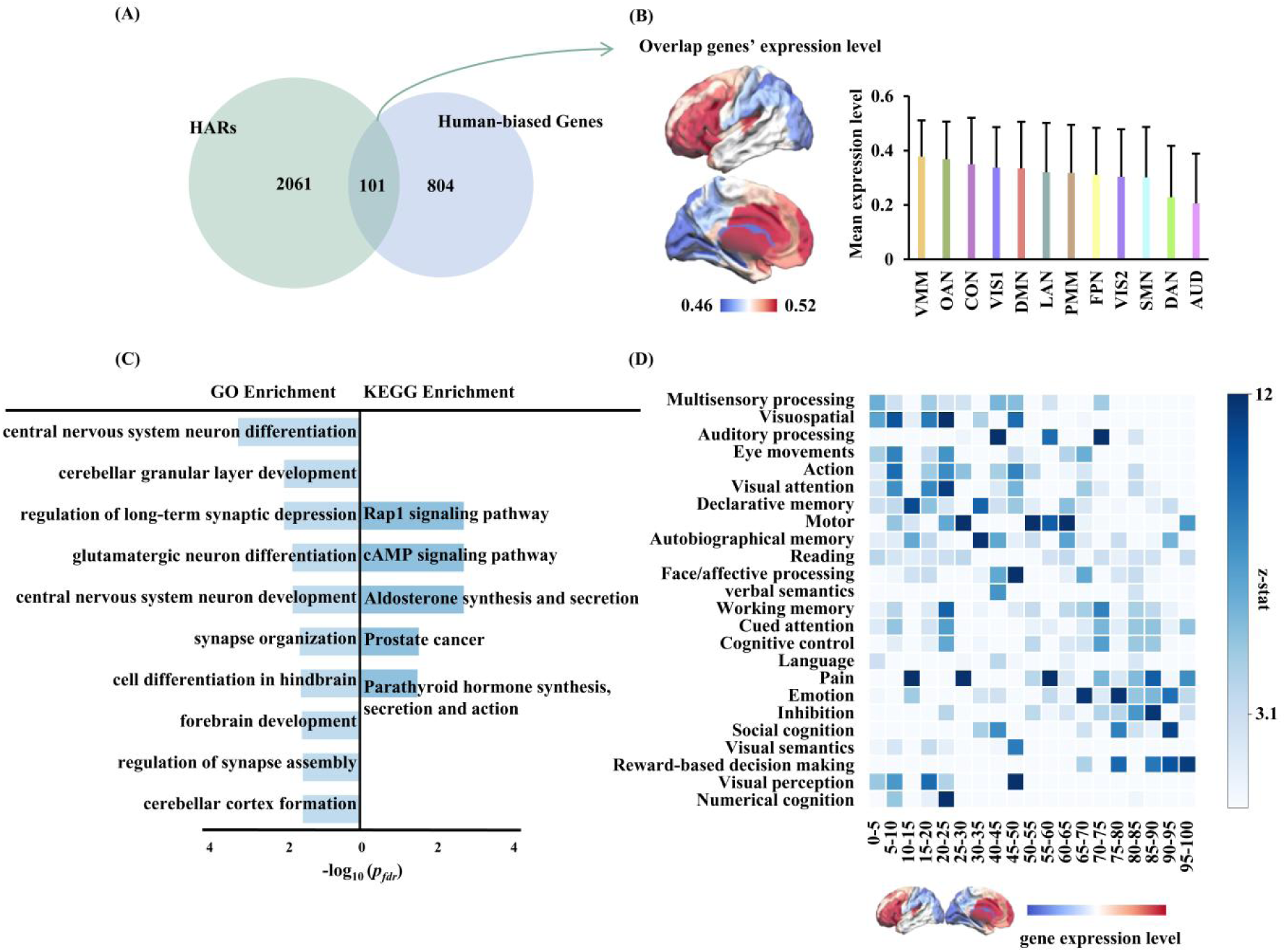
Integrative analysis of human-specific genes associated with SFC and HAR genes. (A) Overlap between 2,162 human-accelerated region (HAR) genes and the 905 human-specific genes significantly linked to SFC, yielding 101 overlapping genes. (B) Spatial expression patterns of these 101 genes mapped across 46 left-hemisphere cortical regions, together with their network-level distributions across the 12 canonical functional networks defined by the CAB-NP atlas. (C) GO and KEGG enrichment analyses of the 101 overlapping genes, highlighting biological processes and signaling pathways enriched in this HAR-linked gene subset. (D) Functional decoding of the cortical expression map derived from the 101 HAR-SFC overlap genes using the NeuroSynth meta-analytic framework revealed associations with 24 cognitive domains. Regions with lower expression were preferentially linked to visuospatial processing and eye-movement components, whereas regions with higher expression aligned with emotion and social cognition. Note: *p_fdr_*: FDR corrected.

## Discussion

In this study, we integrated multimodal MRI and cross-species transcriptomic data to characterize evolutionary differences in SFC between humans and macaques. Both species shared a common organizational backbone, with evolutionarily conserved regions, such as the visual and sensorimotor cortices exhibiting strong coupling that supports stable structure-function alignment essential for basic perceptual and motor processes. In contrast, humans displayed markedly lower coupling in transmodal association areas, particularly within the lateral temporal cortex, indicating enhanced functional flexibility that may support higher-order cognition. Across species, SFC was negatively associated with cortical expansion, suggesting that the evolutionary enlargement of association cortices is accompanied by a relaxation of structural-functional constraints. At the molecular level, macaque SFC was primarily associated with pathways involved in cellular maintenance and homeostasis, whereas human SFC was enriched for genes in cell types of astrocyte and oligodendrocyte related to neurodevelopment, myelination, and synaptic remodeling. Moreover, human-specific SFC was primarily associated with diseases of schizophrenia and Alzheimer’s disease and was modulated by HAR-associated genes highly expressed in the prefrontal and association cortices, indicating that evolutionary innovations in gene regulation may have facilitated the emergence of advanced cognitive abilities. Collectively, these findings highlight distinct spatial and molecular organizational principles of SFC across species, providing empirical insight into the neural and genetic foundations of primate brain evolution.

### Spatial distribution patterns of SFC in the human-macaque brain

Cross-species comparisons revealed that both humans and macaques exhibit high SFC in the visual cortex and mPFC, but the evolutionary implications of these regions differ greatly. The visual cortex is a prototypical evolutionarily conserved region, characterized by stable topological architecture and tightly constrained structure-function correspondence, ensuring high-fidelity perceptual processing (63, 64). In contrast, the medial prefrontal cortex exhibits both functional segregation and strong SFC (5). This finding suggests that the mPFC is highly integrated both structurally and functionally that it maintains dense white matter connections with other core nodes of the default mode network (DMN), such as the posterior cingulate cortex (5). This anatomical foundation enables high functional coordination. High SFC may help the mPFC reduce interference from task-positive networks (e.g., the frontoparietal control network) during task execution, thereby maintaining stable working memory performance, particularly when suppression of internally oriented thoughts (e.g., mind-wandering) is required. Thus, although the mPFC is a transmodal area, it exhibits a tightly coupled and relatively stable structure-function relationship during evolution, aligning with its role as a core hub of the DMN (5).

In contrast, the lateral temporal cortex in humans exhibits relatively low SFC. As an essential hub for multimodal integration involved in language, semantic processing, and social cognition, this region relies less on direct white matter pathways. Instead, its functional connectivity is often mediated by polysynaptic or dynamic functional interactions (65, 66). Consequently, these regions display lower SFC (5, 7), reflecting greater communication flexibility and functional dynamics. Such lower SFC may constitute an important neural substrate enabling humans to achieve enhanced adaptability in complex cognitive domains, such as language and abstract reasoning.

Human brain evolution represents a fundamental trade-off between the stability of networks conserved from primate ancestors and the flexibility afforded by newly expanded and uniquely human cortices. Our cross-species analyses revealed that the whole-brain SFC pattern in humans, as well as the interspecies differences in SFC between humans and macaques, was significantly and negatively associated with evolutionary cortical expansion. Evolutionarily conserved regions, such as the primary visual cortex and mPFC, exhibited higher SFC. Relevant literature indicates that the premotor network controlling eye movements is evolutionarily conserved across primates (67), with its relatively high SFC reflecting a stable structure-function correspondence that supports core functions such as self-referential processing and internally directed cognition (5). This evolutionary stability is also evident in the mPFC, where, despite certain subregions being associated with uniquely human higher-order cognitive functions, the overall functional organization remains highly consistent between humans and macaques, suggesting a notable degree of evolutionary conservation (68, 69). In contrast, regions such as the lateral temporal cortex in humans, particularly Wernicke’s area, have undergone substantial cortical expansion, supporting complex cognitive functions such as language and abstract thinking (5, 70, 71). Overall, these findings suggest that cortical expansion during evolution was accompanied by a reduction in SFC, thereby endowing newly evolved association areas with greater functional flexibility.

Our results indicate that the SFC pattern observed under anesthesia highly reflects that of the awake state, while we also acknowledge the potential influence of anesthesia. Studies have shown that under light anesthesia, the overall organization of SFC remains comparable to that in the awake brain, with functional networks more tightly constrained by underlying anatomical connectivity (29). For instance, Xu et al. compared functional connectivity matrices between awake and anesthetized macaques under monocrystalline iron oxide nanoparticle (MION)-enhanced conditions across multiple centers and found highly similar spatial distributions (72). Under MION conditions, the stability of functional connectivity showed negligible differences between states, suggesting that anesthesia does not disrupt the fundamental architecture of SFC (72). Nevertheless, anesthesia does exert systematic effects: the correlation between structural and functional connectivity tends to increase, while awake brains exhibit richer and more flexible dynamics beyond structural constraints (73). Moreover, functional configurations during anesthesia often show reduced anticorrelations and greater rigidity, reflecting stronger anatomical constraints, whereas the awake state allows the brain to explore a wider range of dynamic functional configurations (74).

### Molecular basis of SFC

The transcriptomic analysis revealed that the spatial distribution patterns of SFC in humans and macaques are regulated by distinct molecular features, reflecting species-specific evolutionary trajectories. In macaques, genes significantly associated with SFC were predominantly enriched in pathways involved in fundamental cellular maintenance processes, including protein homeostasis, RNA processing, intracellular transport, and epigenetic regulation. These pathways are essential for sustaining neuronal survival and stability, suggesting that macaque SFC may primarily reflect the cellular mechanisms required to maintain robust structure-function integration within the sensorimotor system. Further analysis demonstrated that macaque-specific SFC-associated genes were significantly enriched in “post-translational protein modification” pathways (75). A well-orchestrated and precise network of post-translational modifications helps maintain the predictability of neuronal molecular composition and functional states, thereby ensuring consistency between structural substrates (e.g., axons, dendrites, and synapses) and their functional activity (76, 77). This mechanism is particularly important in sensorimotor circuits that demand high-fidelity information transmission and may account for the higher SFC observed in these regions. Moreover, macaque SFC showed strong associations with the “establishment of localization within cells”, whereby proteins and RNAs are accurately transported and localized to specific cellular compartments. Reliable molecular localization strengthens the correspondence between anatomical connectivity and functional activity patterns, thereby enhancing structure-function coupling at local and regional levels (78, 79). Collectively, these findings suggest that the molecular architecture underlying macaque SFC is more oriented toward maintaining neuronal structural stability and functional consistency, rather than promoting large-scale network plasticity, which may be crucial for the precise and efficient signal transmission required by the sensorimotor system.

In contrast, the genes associated with human-specific SFC were mainly expressed in cell types of oligodendrocytes and astrocytes and were predominantly associated with neurodevelopment, myelination, and synaptic regulation. These molecular programs likely facilitate the establishment of highly plastic and efficient long-range connections, which in turn support complex cognitive functions. This observation aligns with previous studies showing that prolonged myelination and synaptic specialization are key features of human cortical evolution, enabling effective communication across distributed association areas (80–82). Among them, oligodendrocytes promote myelination to support efficient long-range signal transmission (83), while astrocytes contribute to synaptic regulation, facilitating effective neuronal communication (84–86). Taken together, the enrichment of human-specific SFC genes may enhance the plasticity and efficiency of long-range connections through prolonged myelination and synaptic refinement, thereby facilitating the evolutionary emergence of complex cognitive abilities. Human-specific SFC genes not only promote long-range synaptic plasticity and myelination but also provide a potential molecular basis for genetic susceptibility to psychiatric disorders. Neuropsychiatric disorders, including schizophrenia and Alzheimer’s disease, and attention deficit hyperactivity disorder (ADHD), show substantial genetic overlap with human-specific neurodevelopmental programs (87). This overlap underscores the notion that such disorders may represent the “cost of evolutionary innovation” (88), whereby the acquisition of enhanced cognitive flexibility and extended lifespan has inevitably increased the risk of neural dysregulation (87). Schizophrenia, in particular, has been proposed as an evolutionary “cost” of developing language and higher cognitive abilities (89), suggesting that the same gene networks underlying complex linguistic circuits and prolonged myelination may also predispose individuals to this disorder (90). Similarly, studies have found that human-specific regulatory sequences contributing to neocortical expansion carry risk variants associated with Alzheimer’s disease (91).

Importantly, by comparing with HARs genes, we identified the HARs genes that specifically regulate human SFC, which highly expressed in prefrontal and association cortices, particularly concentrated in networks such as the limbic and frontoparietal systems that are critical for higher-order cognition (92, 93). This provides a potential genetic basis for the expansion and functional specialization of these cross-modal regions in humans (5). The enrichment of HAR-related genes specifically regulating human SFC revealed that these genes mainly involved in synaptic plasticity, axon guidance, and intracellular signal transduction suggesting that innovations in regulatory elements during evolution may have optimized the architecture of SFC to support uniquely human abilities, including working memory, language, and social cognition.

Despite our considerable efforts to minimize possible biases, several constraints must be taken into account when interpreting these findings. First, the neuroimaging data of macaques were acquired under anesthesia. Although analyses of an independent awake macaque dataset demonstrated that whole-brain FC and SFC patterns closely resembled those observed under anesthesia, we acknowledge that anesthesia may still alter subtle neural dynamics and coupling features. Future studies should prioritize collecting large-scale awake macaque datasets to validate the present findings. Second, the transcriptomic data were derived from public atlases rather than the same individuals used for imaging. Although we used cross-species shared genes, rigorous preprocessing, and spatial permutation controls to mitigate potential bias, confirming causal relationships between imaging phenotypes and gene expression will require acquiring both modalities from the same cohort. Third, the sample size was relatively limited, which may reduce statistical power and generalizability. Finally, the transcriptomic data were based on bulk regional expression, limiting the ability to capture cell-type-specific or spatiotemporal heterogeneity underlying SFC. Although our enrichment analyses identified pathways associated with distinct cell types, future studies combining cross-species single-cell or single-nucleus transcriptomic atlases with imaging parcellations will help clarify the cellular basis of interregional SFC differences.

## Conclusion

This study provides a cross-species framework to elucidate how SFC has evolved across the primate lineage. By integrating multimodal MRI with transcriptomic data, we demonstrate that while core sensorimotor regions remain evolutionarily conserved and tightly coupled, the association cortices-particularly in humans-exhibit weaker coupling, supporting flexible integration essential for higher cognitive functions. At the molecular level, the divergence of SFC reflects distinct genetic regulation: in macaques, coupling is anchored in conserved processes that maintain structural stability, whereas in humans, it aligns with genes involved in neurodevelopment, myelination, synaptic plasticity, and HARs, suggesting an evolutionary shift toward molecular mechanisms that enable cognitive innovation. Collectively, these findings uncover a dual evolutionary strategy that balances structural conservation with functional adaptability, offering novel insights into the molecular and network-level mechanisms underlying the emergence of human cognitive specialization.

## Data and code availability statement

The data and code that support the findings of this study are available on request from the corresponding author.

## Acknowledgement

This study was supported by the Yunnan Fundamental Research Projects (202501AV070005).

## Author contributions

Jiaojian Wang contributed to the conception and design of the study; Jianing Ma, Wei Li, Yingzi Ma, Jin Chen, Jing Su, Yashi Wu, and Chengsi Luo contributed to the acquisition of data and data analysis; Jianing Ma and Jiaojian Wang contributed to the drafting and editing of the manuscript; Jiaojian Wang, Jianing Ma, and Wen Li contributed to the critical review of the manuscript. All authors commented and discussed the results.

## Compliance with ethical standards

All experimental procedures were conducted in strict compliance with national guidelines and regulations on animal protection and use, and the protocols were approved by the National Bureau of Animal Research of China and the Animal Care Committee (permit number: KUST202301018 and KMUST-MEC-207).

