## Supplementary Materials for "Evolutionary Divergence of Structure-Function Coupling between Human and Macaque: Spatial Patterns and Transcriptomic Basis"

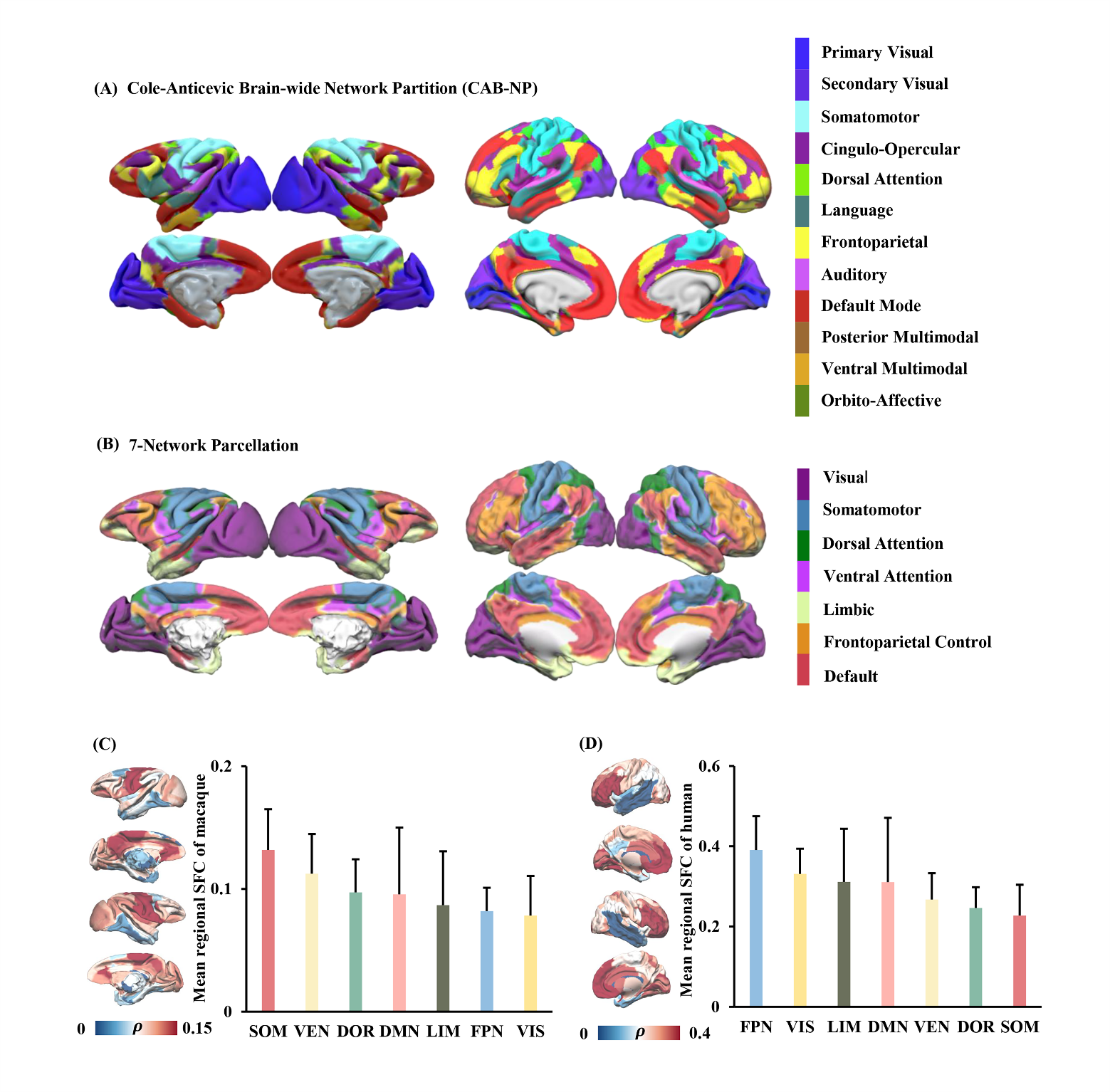
**Figure S1. Cross-species functional network parcellations and SFC distribution.** (A) Spatial layout of the 12 functional networks from the CAB-NP displayed on the human cortical surface and projected onto the macaque cortex using the macaque-to-human spherical deformation field. (B) Spatial distribution of the canonical 7-network parcellation in humans and macaques, used to evaluate the robustness of SFC patterns across alternative parcellation schemes. (C) Mean SFC values averaged within each of the seven canonical functional networks for humans and macaques.

**Figure S2. Human-macaque differences in SFC.** (A) Brain regions exhibiting significant differences in SFC between humans and macaques, identified using two-sample *t*-tests with Bonferroni correction (*p* < 0.05). (B) Visualization of regions with positive and negative *t*-values. Positive *t*-values indicate higher SFC in humans, with the strongest effect observed in the medial prefrontal cortex, whereas negative *t*-values indicate higher SFC in macaques, with the lowest *t*-value observed in the superior temporal cortex. The table on the left lists all regions showing significantly positive and negative *t*-values.

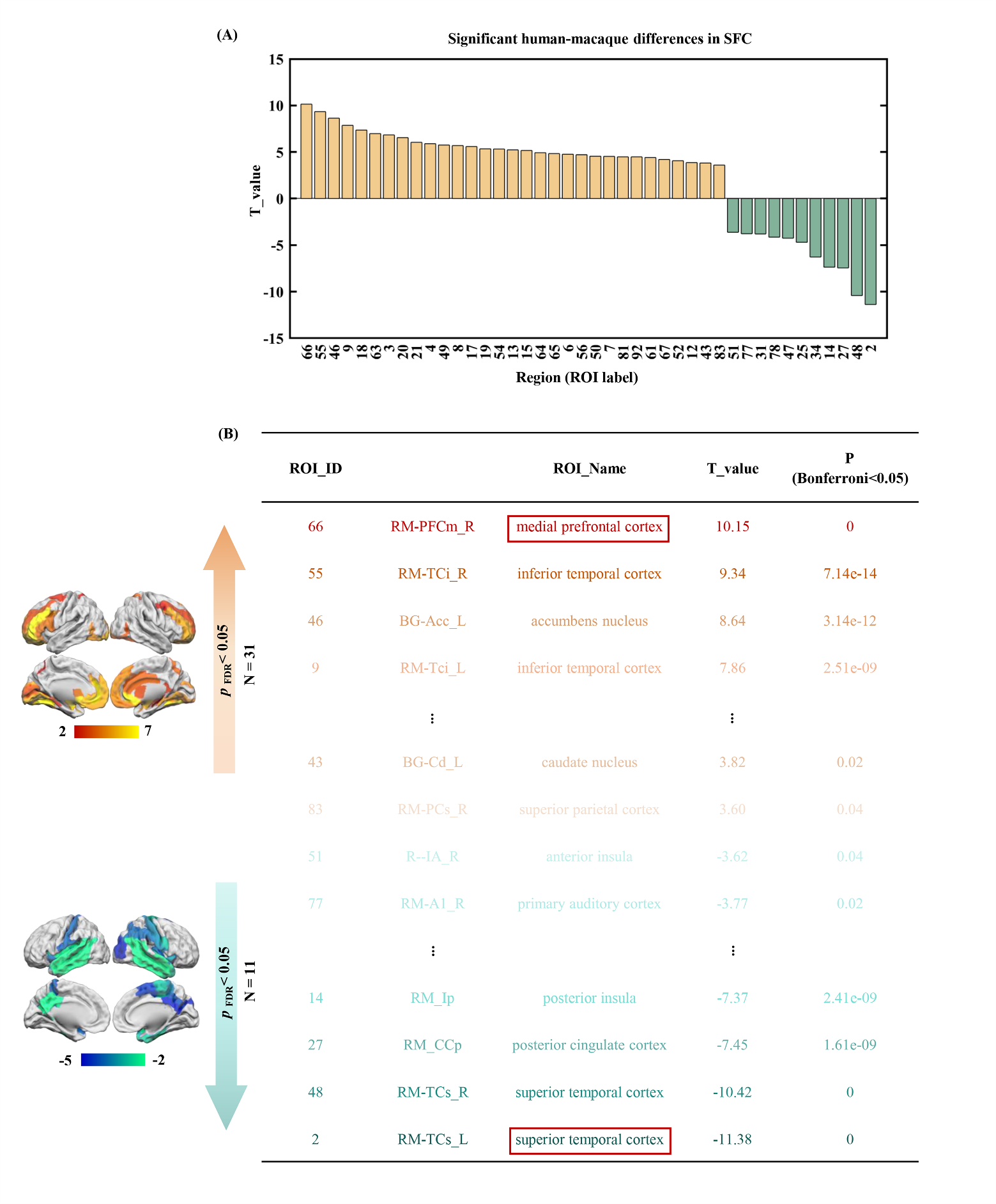
**Figure S3. Functional decoding of human SFC.** Associations between human SFC values and 24 cognitive domains derived from the NeuroSynth meta-analytic database. Cognitive domains are shown as rows and are sorted by the weighted mean of their *z*-statistic values. The results showed that regions with lower SFC in humans were primarily associated with emotion and declarative memory, whereas regions with higher SFC were linked to motor and
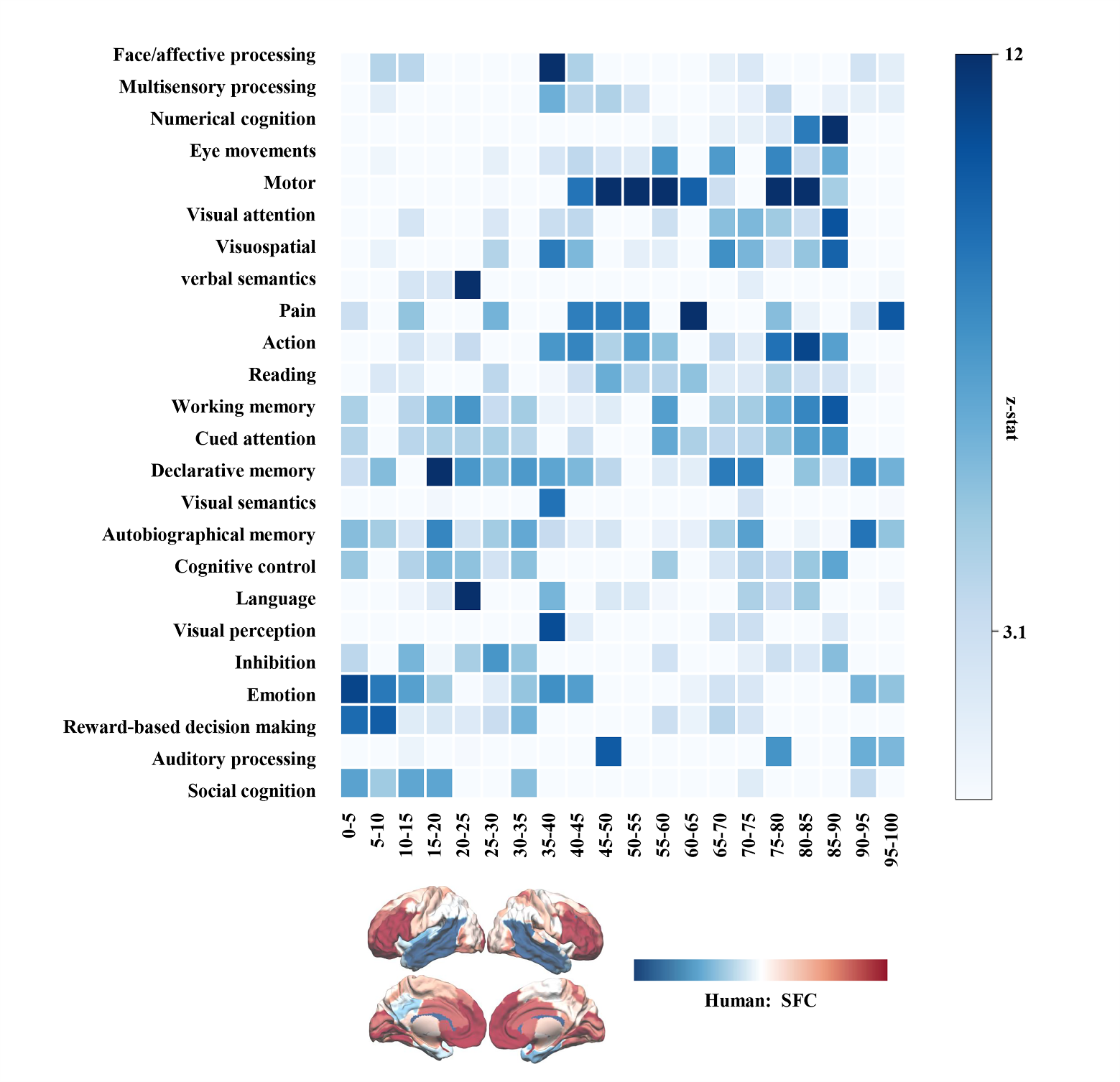
visuospatial functions.

**Figure S4. Validation of SFC using deterministic tractography in macaques.** SFC derived from probabilistic tractography was validated against structural networks generated by deterministic tractography. Significant correlations were found for SFC based on the number of streamlines (NUM; *r* = 0.75, *p* < 0.001), mean diffusivity (MD; *r* = 0.46, *p* < 0.001), and fractional anisotropy (FA; *r* = 0.23, *p* < 0.005).

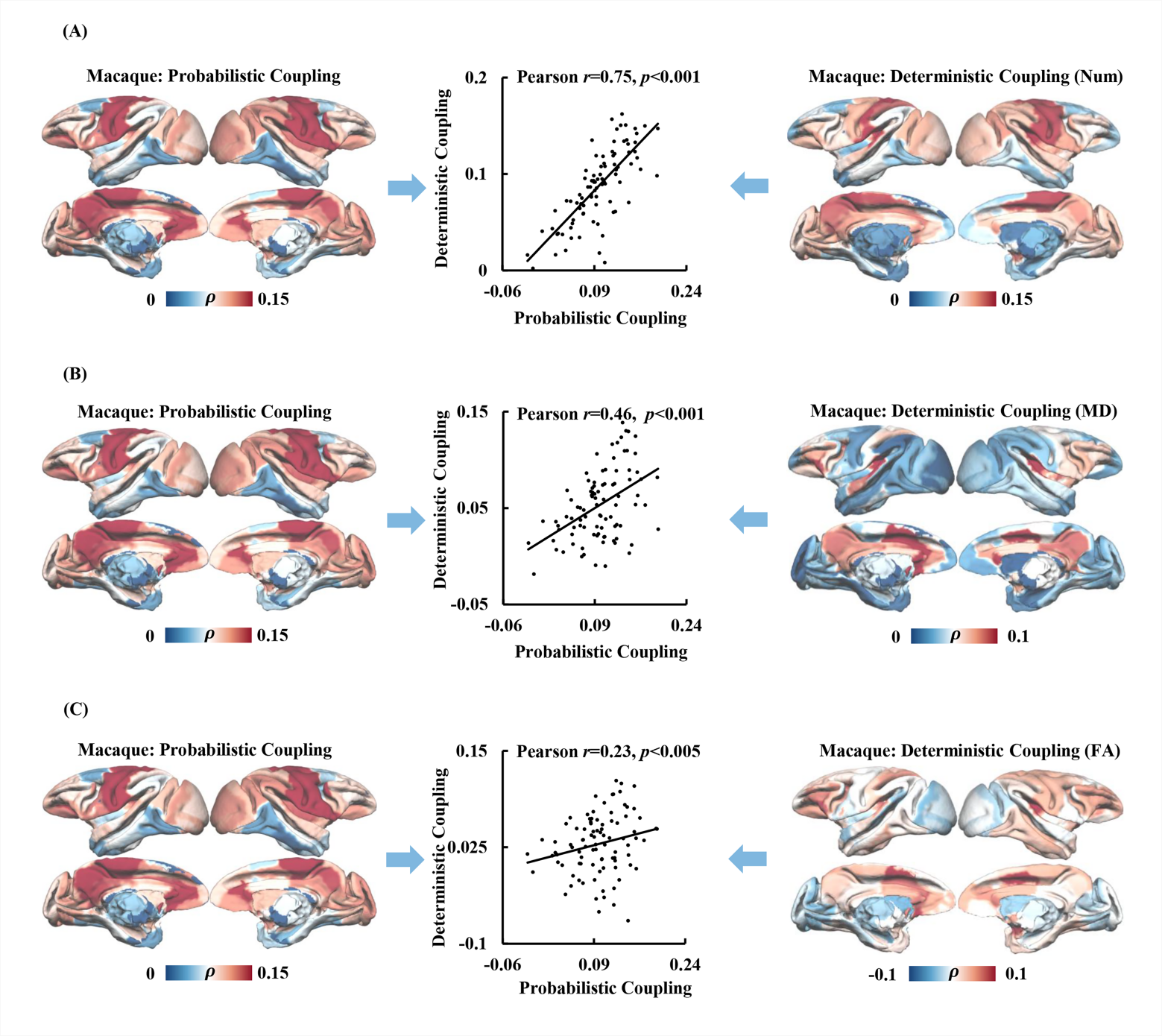

**Figure S5. Validation of SFC using deterministic tractography in humans.** SFC derived from probabilistic tractography was validated against structural networks generated by deterministic tractography. Significant correlations were observed for SFC based on the NUM (*r* = 0.65, *p* < 0.001), MD (*r* = 0.56, *p* < 0.001), and FA (*r* = 0.41, *p* < 0.001).

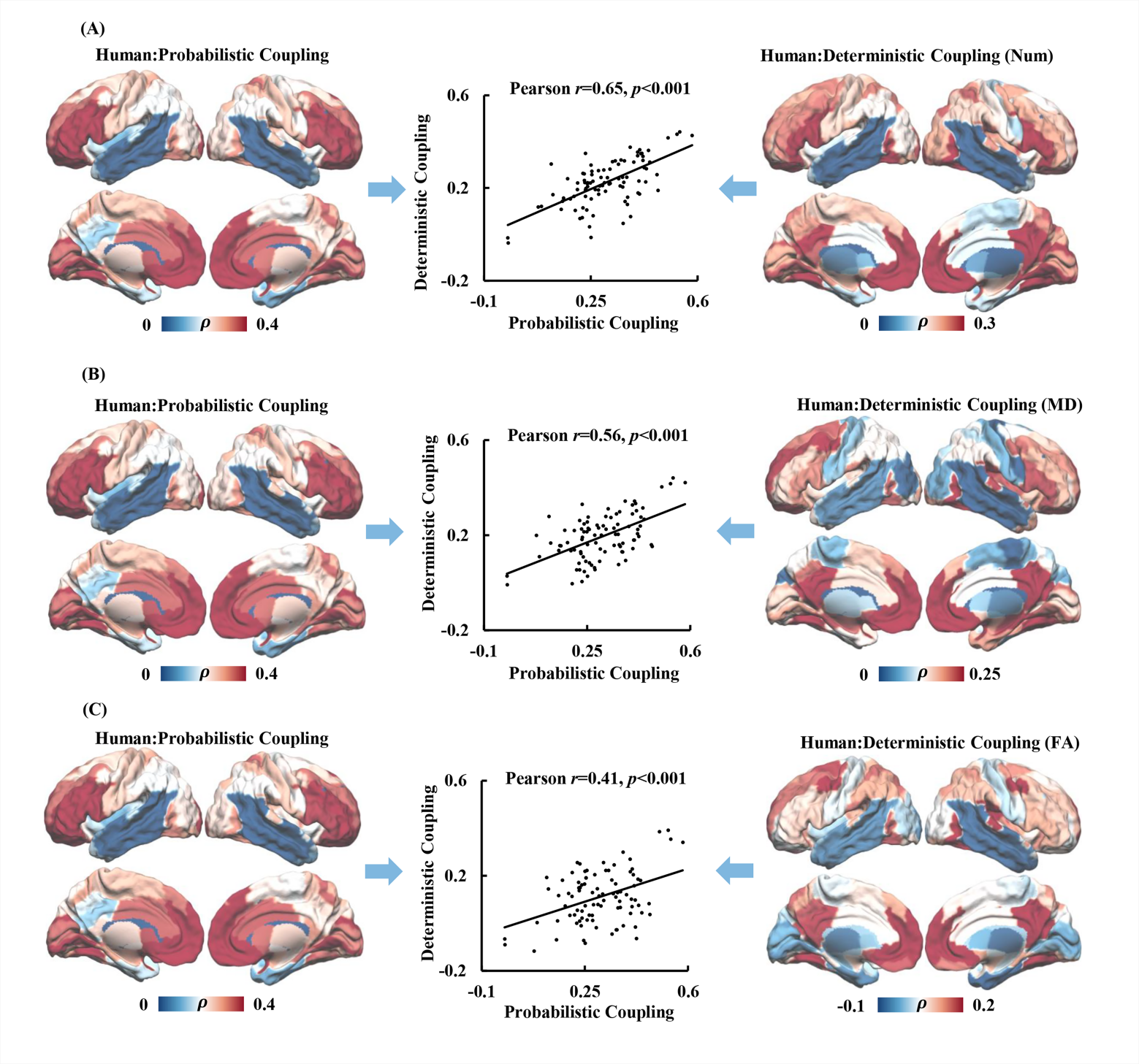

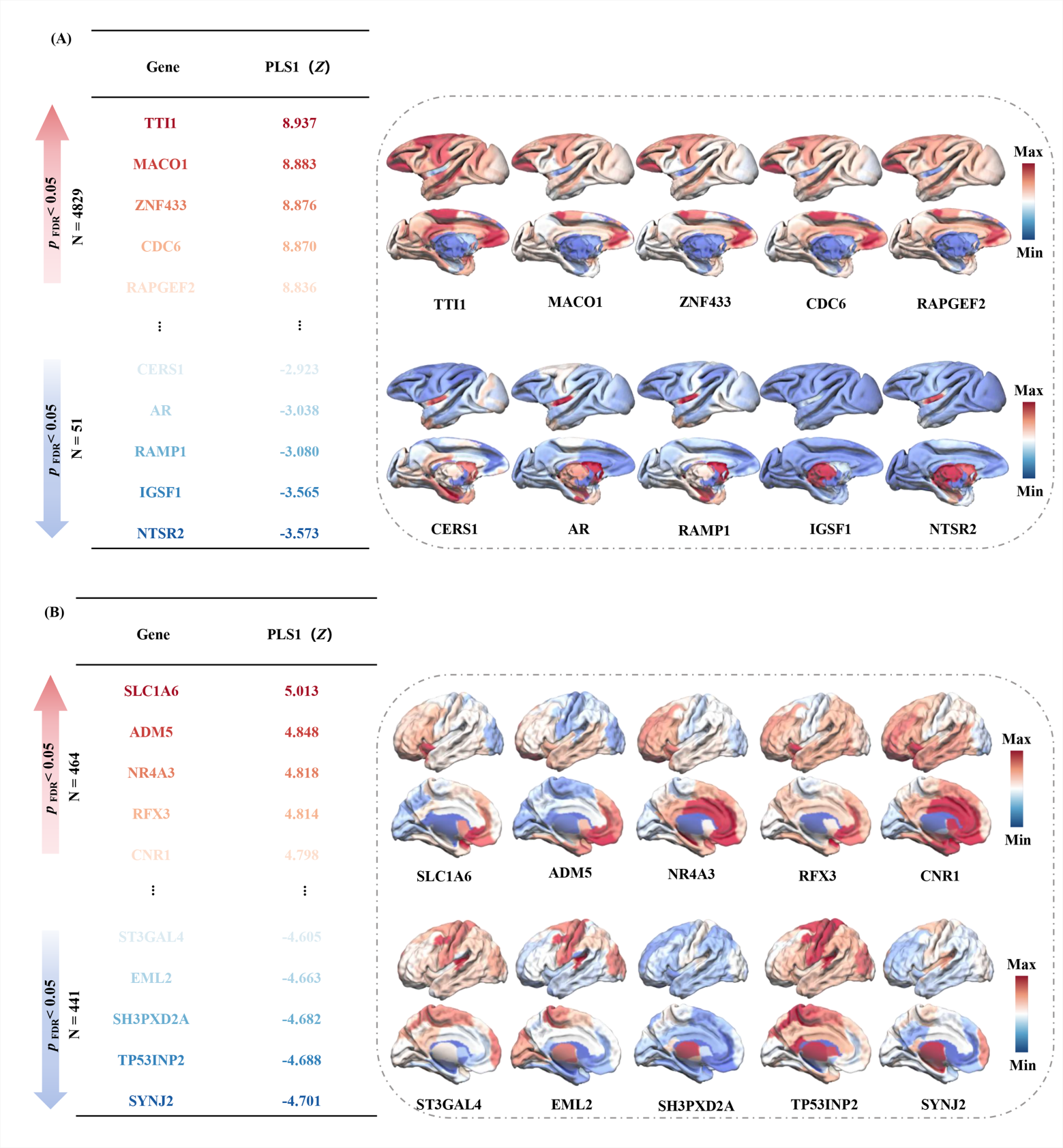
**Figure S6. Significant positive and negative genes identified from PLS1 and their cortical expression patterns.** (A) The top five positively weighted and top five negatively weighted genes (FDR *p* < 0.05) were identified from the first partial least squares component (PLS1) linking macaque SFC with regional gene expression. Spatial expression maps for each gene are shown across the macaque cortex. (B) The same analysis was performed in humans, displaying the top five positively and negatively weighted genes along with their cortical expression maps.

**Figure S7. Functional decoding of left-hemisphere gene expression significantly associated with human SFC.** The analysis examined associations between the left-hemisphere gene expression significantly associated with human SFC and 24 cognitive domains derived from the NeuroSynth meta-analytic database. The decoding results indicated that regions with lower mean expression were primarily related to visuospatial processing, whereas higher expression was linked to reward-based decision making, social cognition, and emotion.

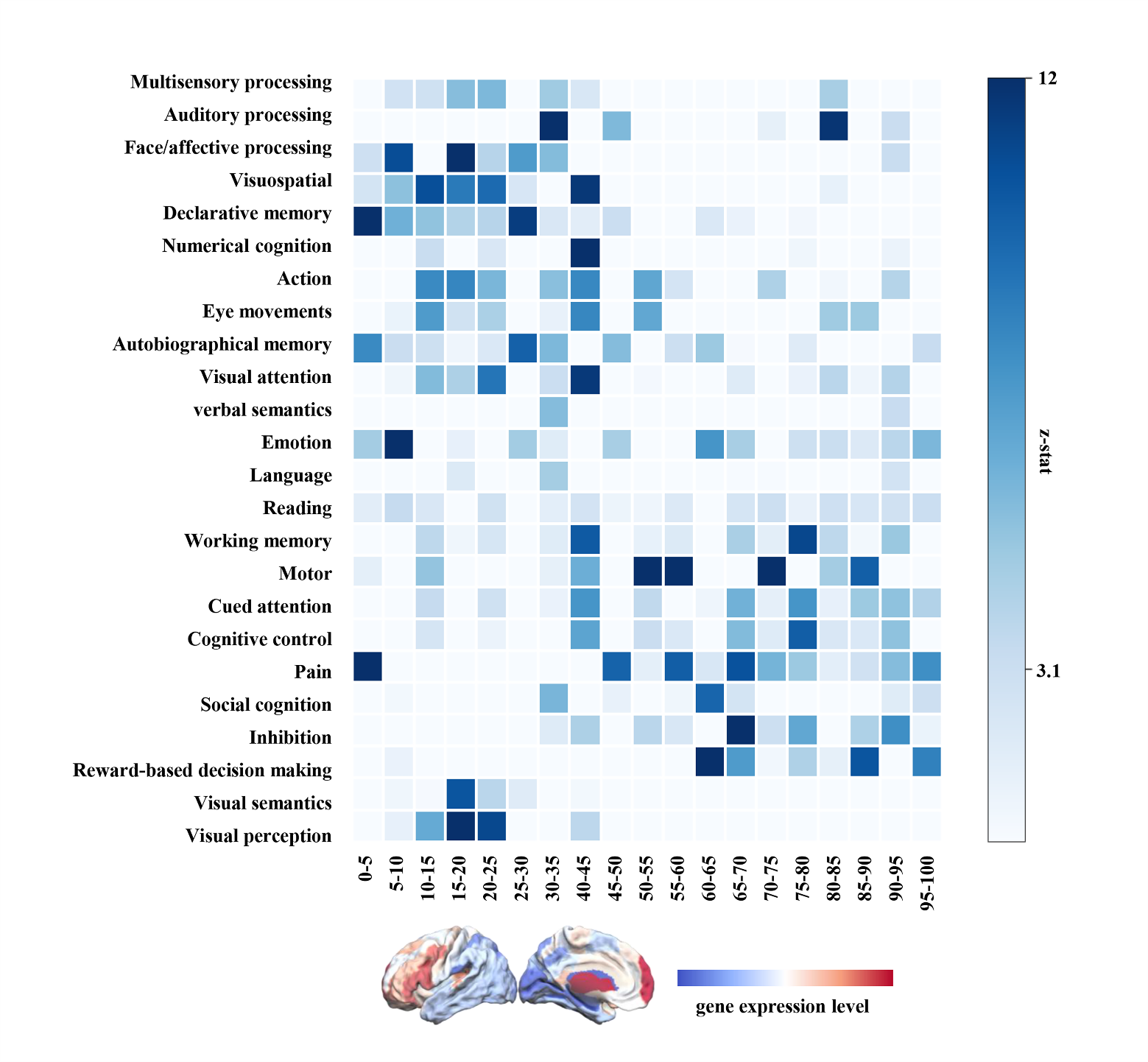

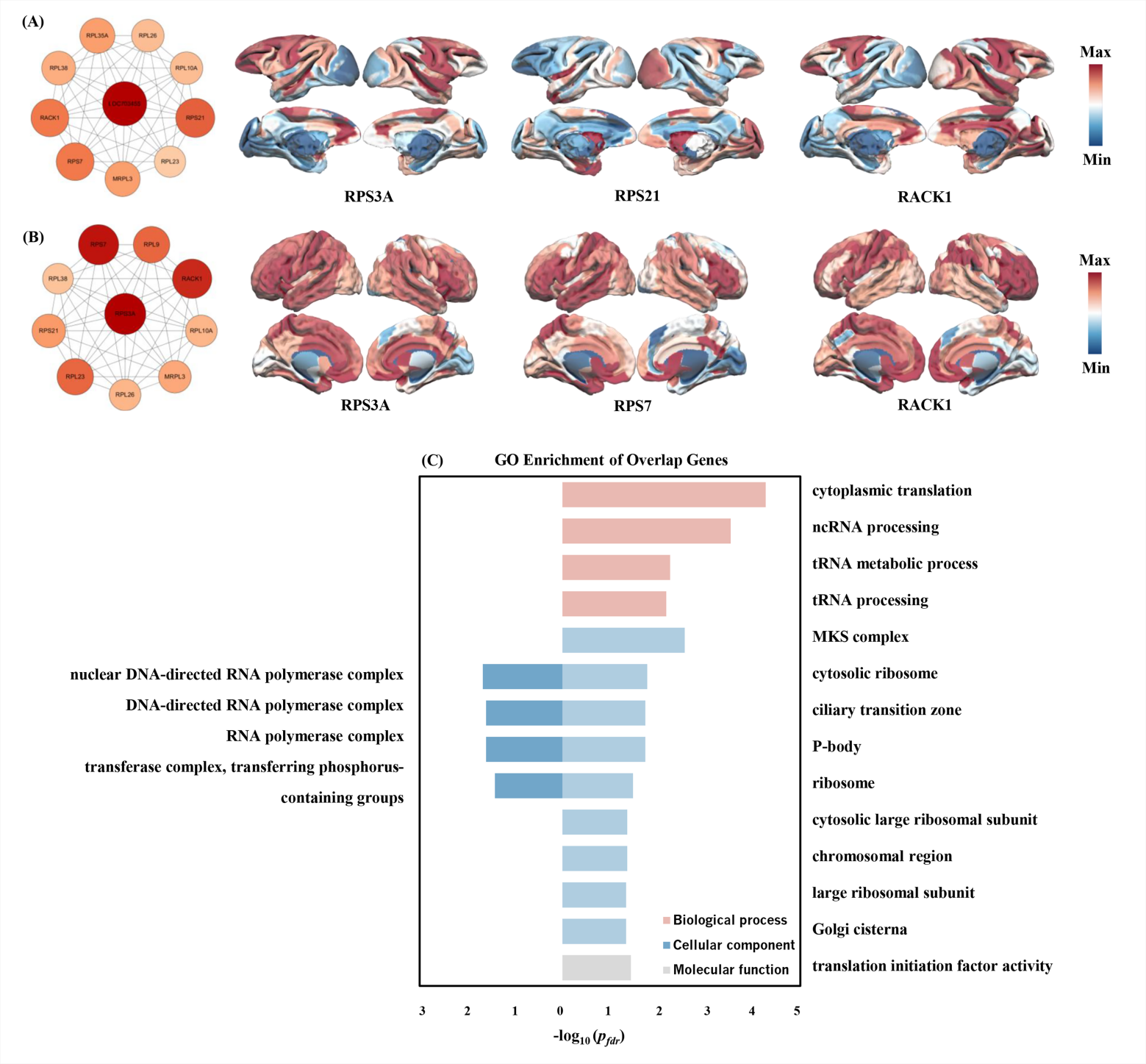
**Figure S8. Protein-protein interaction and functional enrichment of human-macaque overlap genes.** (A) PPI network of the 818 human-macaque overlap genes constructed using the human gene background, highlighting the top three hub genes ranked by node degree and their whole-brain expression patterns. (B) PPI network of the same 818 overlap genes in the macaque gene background, showing the three highest-degree hub genes and their cortical expression maps. (C) GO enrichment analyses for the overlap genes, presented separately for macaques (left) and humans (right).
